# Autologous host-pathogen pairing enhances early immune responses and bacterial growth control in tuberculosis *in vitro* granuloma model

**DOI:** 10.1101/2024.12.20.629565

**Authors:** Charlotte Genestet, Chloé Bourg, Elisabeth Hodille, Olivier Bahuaud, Florence Ader, Oana Dumitrescu Lyon TB study group

## Abstract

Tuberculosis (TB), caused by *Mycobacterium tuberculosis* (Mtb), remains a global health challenge, characterized by significant heterogeneity in its clinical presentation and physiopathology, underscoring the need for optimized models to better understand host-pathogen interactions. This study evaluated the impact of both host and Mtb clinical isolate variability on bacterial growth and dormancy and immune responses using an *in vitro* granuloma model. Peripheral blood mononuclear cells from active TB patients were infected with autologous and heterologous Mtb clinical isolates, as well as the H37Rv Mtb reference strain. Bacterial growth and dormancy dynamics and cytokines release were assessed at 0-, 7-, 14- and 21-days post-infection. Results revealed that bacterial growth and dormancy dynamics were mostly isolate-dependent, with almost no impact of PBMC donor. However, at 7-days post-infection, autologous infections exhibited lower bacterial loads and elevated Th1-type cytokines, such as TNF-α and IFN-γ. Yet, this early advantage in bacterial control did not persist at later time points. These findings highlight the importance of host-pathogen pairing in shaping immune responses and validate the *in vitro* granuloma model as a relevant tool for studying TB pathogenesis and evaluating therapeutic interventions. Furthermore, further research is needed to decipher the specificities of autologous immune response.

## INTRODUCTION

Tuberculosis (TB) remains one of the leading global health threats, causing nearly 4,000 deaths daily (WHO, 2023). TB is caused by *Mycobacterium tuberculosis* (Mtb), that primarily targets the lungs, leading to the formation of complex immune structures known as granulomas—a hallmark of Mtb infection. Granulomas are organized aggregates of differentiated macrophages surrounded by lymphocytes that contain Mtb and restrict its growth. However, granulomas also provide a microenvironment in which Mtb can persist, impeding TB treatment and eradication. Studying these granulomas is essential for understanding TB pathogenesis and developing effective treatments.

In recent years, *in vitro* granuloma models have emerged as valuable tools for TB research, replicating the structure and cellular environment of granulomas under controlled conditions (Elkington et al., 2019; Ganesan et al., 2022; Puissegur et al., 2004). These models are formed by infecting peripheral blood mononuclear cells (PBMCs) with Mtb, leading to the spontaneous formation of granuloma-like structures. However, variables within these models, such as the genetic variability of both the host and the pathogen, remain underexplored. Previous studies have focused on either donor variability (e.g., healthy versus latent TB patients) or Mtb strain variability (e.g., clinical isolates) but none both simultaneously. Yet, various approaches indicate significant inter-individual variability in susceptibility to Mtb infection (Azad et al., 2020; Musvosvi et al., 2023; Sadee et al., 2023; Simmons et al., 2018). Furthermore, while the Mtb complex is clonally derived, ten major lineages of human-adapted Mtb have been identified, differing in intracellular survival and immune response induction (Arbués et al., 2024; Guyeux et al., 2024; Homolka et al., 2010; Mourik et al., 2019; Wang et al., 2010). Recent studies using macrophage infection models confirm that host-pathogen pairing influences Mtb uptake and growth (Gröschel et al., 2023; Osei-Wusu et al., 2023).

These findings highlight the need for models addressing both host and pathogen variability to better understand TB pathogenesis. Accordingly, in this proof-of-concept study, we used PBMCs from TB-treated patients infected with their autologous Mtb clinical isolates, heterologous isolates, and the H37Rv reference strain. The dynamics of Mtb growth and dormancy, as well as cytokine response patterns, were evaluated to investigate the impact of host-pathogen pairing in the *in vitro* granuloma model.

## RESULTS

### Donors and clinical Mtb clinical isolates

Experiments were conducted using PBMCs from three patients treated for active TB, referred as P1, P2 and P3. Each patient was infected with a Mtb clinical isolate from a different lineage— lineages 1, 4, and 3 (L1, L4, and L3), respectively. Blood samples were collected at the end of anti-TB treatment. For *in vitro* granuloma formation (Figure S1), patients’ PBMCs were infected with their autologous Mtb clinical isolate or with the Mtb clinical isolates from the other two patients. Additionally, an epidemic Mtb clinical isolate of lineage 2 (L2) responsible of a large regional cluster (Genestet et al., 2019) and the H37Rv reference strain were also used for PBMCs infections for comparison.

### Mtb isolate-dependent bacterial growth dynamics but with improved early infection control in autologous pairings

Bacterial load was assessed by CFU counting after granuloma disruption and cell lysis at 7-, 14-, and 21-days post-infection (dpi). Additionally, Mtb dormancy was evaluated by quantifying differentially culturable bacteria in the presence or absence of resuscitation-promoting factors (Figures 1, S2, S3, S4).

**Figure 1:**
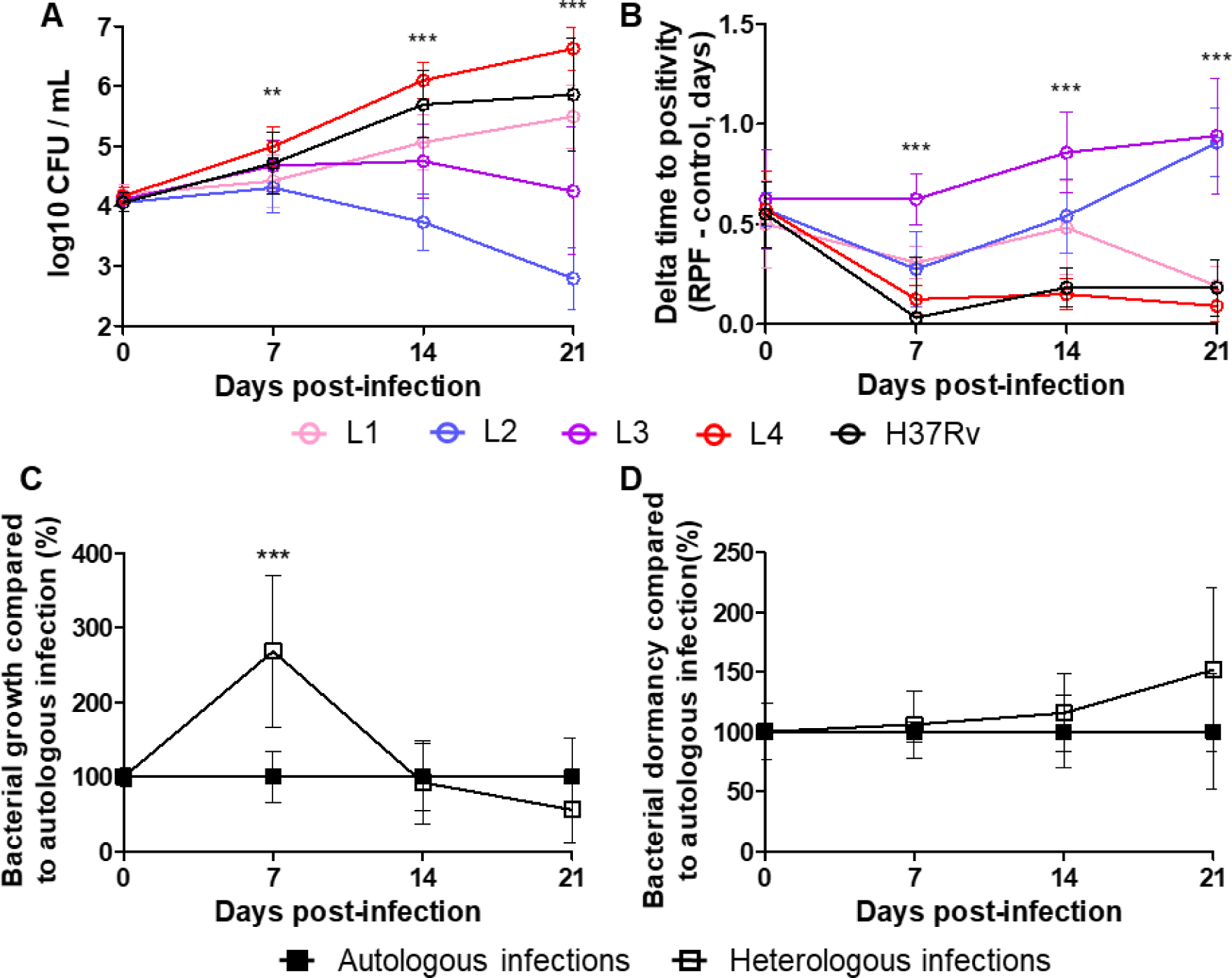
Mtb isolate-dependent bacterial growth and dormancy dynamics with improved early infection control in autologous pairings. PBMC from 3 active TB patients were collected at the end of anti-TB treatment. Each patient was infected with a Mtb clinical isolate from a different lineage—lineages 1, 4, and 3 [L1 (pink circle), L4 (red circle), and L3 (purple circle)], respectively. For *in vitro* granuloma formation, these PBMCs were infected with their own autologous Mtb clinical isolate (full square) as well as with the Mtb clinical isolates from the other two patients (empty square). Additionally, a lineage 2 [L2 (blue circle)] Mtb clinical isolate from a large regional cluster and the H37Rv reference strain (black circle) were included for comparison. (A, C) Bacterial load was assessed by CFU counting after granuloma disruption and cell lysis at 7-, 14-, and 21-days post-infection. (B, D) At the same time-points, Mtb dormancy was evaluated by quantifying differentially culturable bacteria in the presence or absence of resuscitation-promoting factors (RPF) using MGIT time to positivity (TTP) system, reflecting the bacterial growth. (C, D) For each donor, the results obtained were normalized to those of autologous infection. Values for each condition are the mean ± SD of two independent experiments, performed in duplicate. Means were compared at each time-points using Repeated Measures ANOVA test (A, B) or paired t-test (C, D).

Distinct bacterial growth patterns were observed, predominantly influenced by the Mtb clinical isolate rather than by donor. As early as 7-dpi, the L4 isolate exhibited a higher bacterial load than the other isolates, a trend that became more pronounced over time, reaching a load of 6.6±0.36 log_10_ CFU/mL at 21-dpi. The L1 isolate and the H37Rv reference strain showed a progressive increase in bacterial load over time, up to loads of 5.5±0.5 log_10_ CFU/mL and 5.9 ± 0.9 log_10_ CFU/mL at 21-dpi, respectively. The L3 isolate displayed a transient rise in bacterial load at 7-dpi, followed by stasis at 14-dpi and a decline at 21-dpi from 4.7±0.6 log_10_ CFU/mL to 4.2±1.1 log_10_ CFU/mL. In contrast, the L2 isolate exhibited a slight initial increase of the bacterial load at 7-dpi followed by a marked reduction at later time points, with a decline reaching 2.8±0.5 log_10_ CFU/mL (Figure 1A, Figure S2A). The observed reduction in bacterial load for L2 and L3 isolates was associated with an increase in dormancy, as shown by the rise in differentially culturable bacteria in the presence of resuscitation promoting factors, with a growth time delta up to 0.9±0.2 days. Conversely, isolates L1, L4, and the H37Rv reference strain, which had continued bacterial growth, showed significantly lower dormancy induction (Figure 1B and S2B).

When analyzing bacterial growth and dormancy based on donor-specific responses, a notable trend emerged for autologous infections. Specifically, at the early stage of infection (7-dpi), heterologous infections exhibited a higher bacterial load compared to infections using cells autologous to the Mtb clinical isolates, with an increase of 268%±102% (Figure 1C and S3). This finding suggests that early-stage bacterial control may be more effective when host cells are matched to their corresponding clinical isolate. Despite this initial reduction, early bacterial load differences did not translate into improved infection control at later time points and no significant difference was observed in dormancy induction (Figure 1D and S4).

### Mtb isolate-dependent cytokine production dynamics but with enhanced early pro-inflammatory response in autologous pairing

To investigate immune response dynamics, cytokine levels were measured in supernatants collected at each time point. Overall, infections induced a robust Th1-type pro-inflammatory response at early time points, marked by peaks in TNF-α, IFN-γ, and IL-2 alongside notable IL-10 production. The IFN-γ/IL-10 ratio indicated a strong pro-inflammatory response in the initial infection phase. This Th1 response was followed by increased levels of IL-13 and GM-CSF, suggesting a shift toward a Th2-type response and associated macrophage activation (Figure 2).

**Figure 2:**
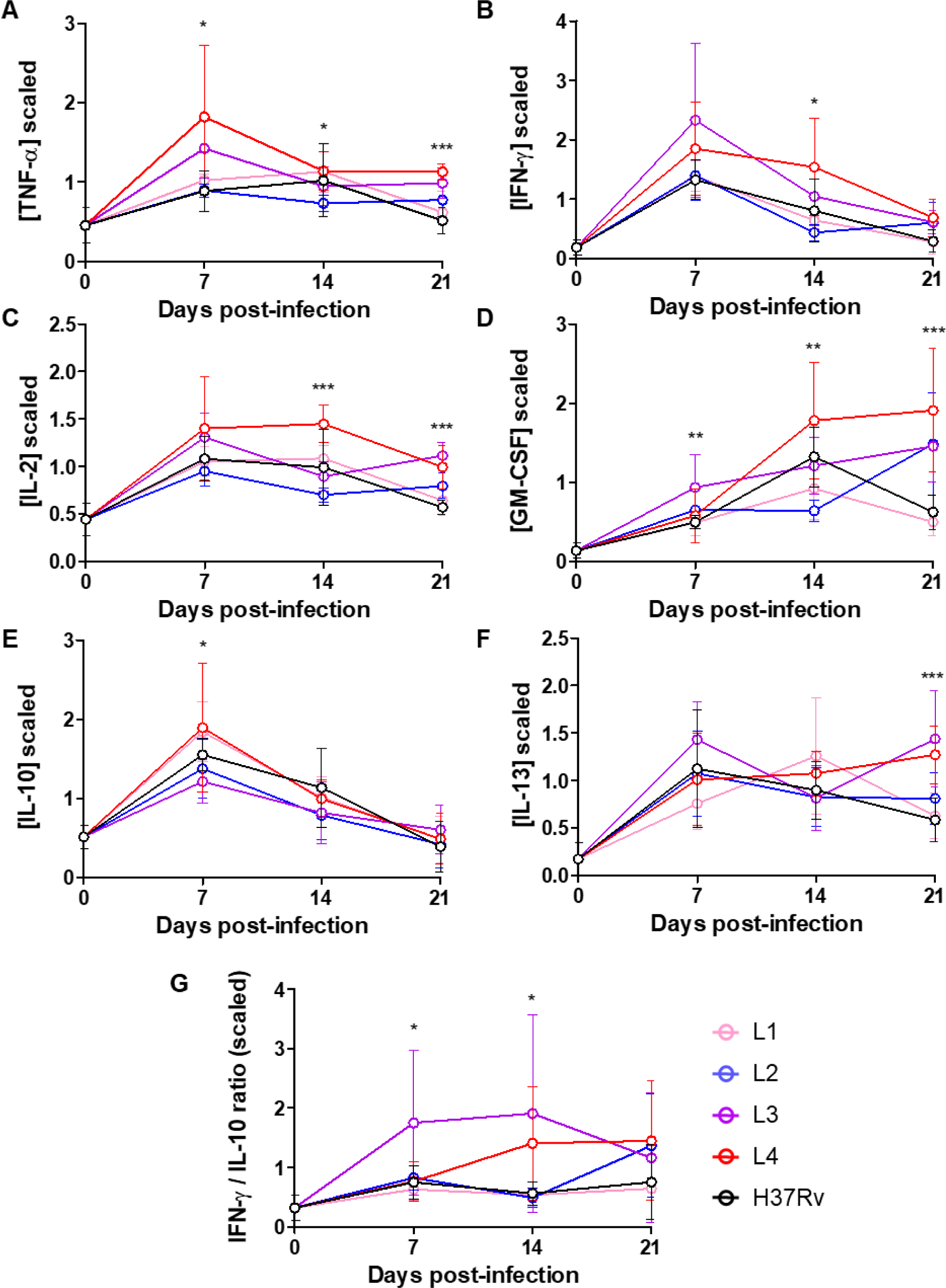
Mtb isolate-dependent cytokine production dynamics during infection. PBMC from 3 active TB patients were collected at the end of anti-TB treatment. For *in vitro* granuloma formation, these PBMCs were infected with their own autologous Mtb clinical isolate as well as with the Mtb clinical isolates from the other two patients, belong to the lineage 1 (pink), lineage 3 (purple) and lineage 4 (red). Additionally, a lineage 2 [L2 (blue)] Mtb clinical isolate from a large regional cluster and the H37Rv reference strain (black) were included for comparison. (A) TNF-α, (B) IFN-γ, (C) IL-2, (D), GM-CSF (E) IL-10, (F) IL-13 release in cell culture supernatant was evaluated by Luminex® Multiplex assay at 7-, 14- and 21-days post-infection. (G) At each time-point IFN-γ/IL-10 ratio was calculated. Values for each condition are the mean ± SD of two independent experiments. Means were compared at each time-points using Repeated Measures ANOVA test.

Mtb isolate-specific analyses revealed marked differences in the timing and intensity of cytokine production (Figures 2 and S5). The L3 isolate induced a rapid and strong Th1-type response, with early cytokine peaks at 7-dpi, which was associated with stabilization of bacterial load (Figure 1). In contrast, L4 isolate triggered a sustained and prolonged pro-inflammatory response, with cytokine peaks between 7- and 14-dpi and continued high pro-inflammatory levels up to 21-dpi (Figure 2), which was associated with a loss of bacterial burden control (Figure 1). In comparison, the L1 isolate and the H37Rv reference strain induced a moderate, progressive inflammatory response aligning with gradual increases in bacterial load. Lastly, the L2 isolate elicited the weakest cytokine response, consistent with low bacterial load (Figures 1 and 2).

Significant differences in cytokine responses were observed when comparing autologous *versus* heterologous infections, particularly in the early infection phase (Figures 3, S6 and S7). Infection of heterologous cells led to a weaker Th1-type response, as evidenced by TNF-α and IFN-γ levels and IFN-γ/IL-10 ratio at 7-dpi which were of 68%±21%, 57%±21% and 47%±14% compared to autologous infections, which correlated with higher early bacterial growth (Figure 1). In contrast, for the L2 isolate and the H37Rv reference strain, cytokine responses were more uniform across donors (Figure S6 and S7). Interestingly, Mtb growth rate was inversely correlated to pro-inflammatory response only for L1, L3 and L4 isolates for which autologous and heterologous associations were tested (Figure S8). This result suggests that the matching of host cells to their corresponding Mtb clinical isolates may enhance initial immune activation and bacterial containment.

**Figure 3:**
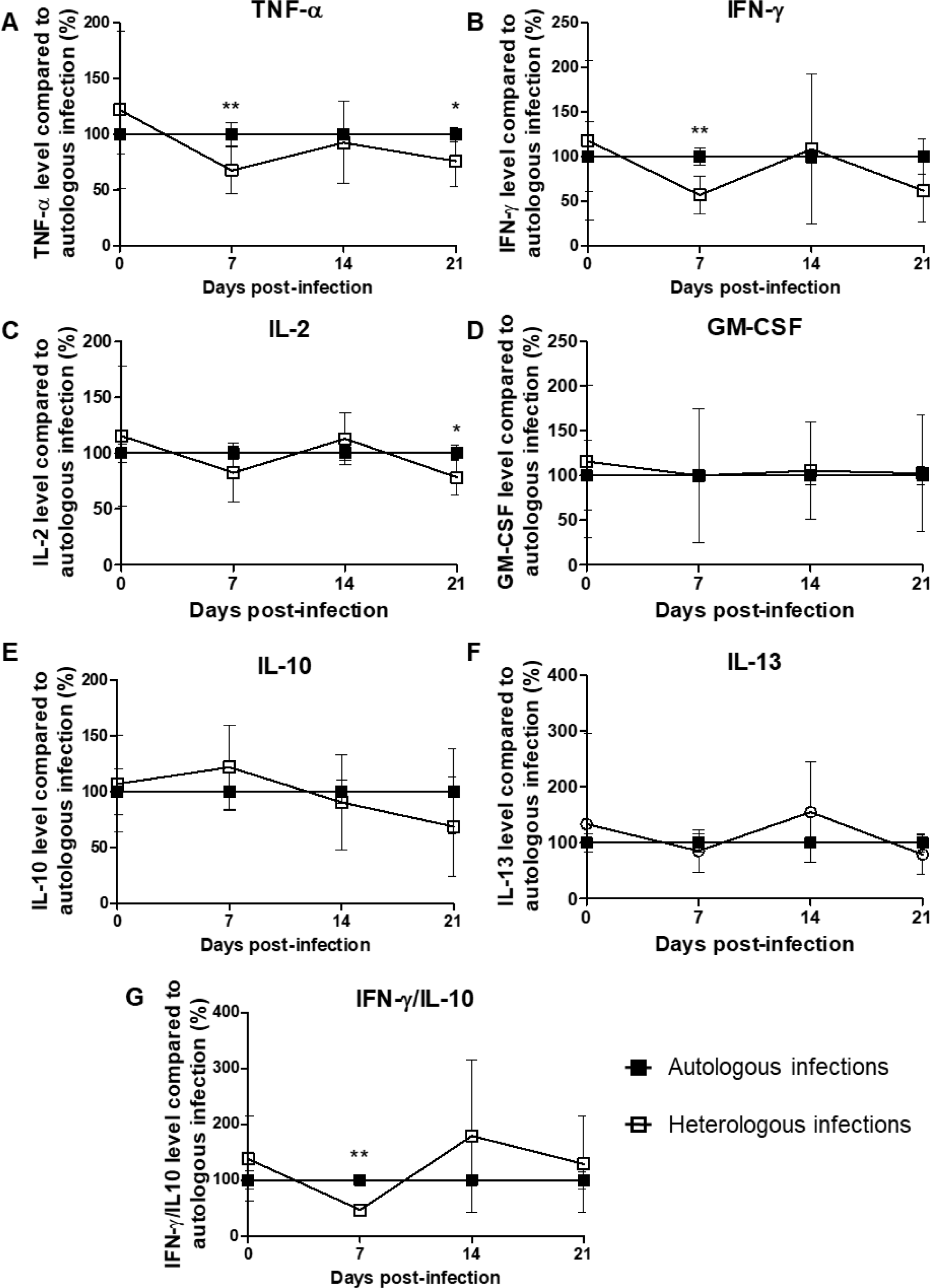
Enhanced early pro-inflammatory response in autologous infections. PBMC from 3 active TB patients were collected at the end of TB treatment. For *in vitro* granuloma formation, these PBMCs were infected with their own autologous Mtb isolate (full square) as well as with the isolates from the other two patients (empty square), belong to the lineage 1, lineage 3 and lineage 4. (A) TNF-α, (B) IFN-γ, (C) IL-2, (D), GM-CSF (E) IL-10, (F) IL-13 release in cell culture supernatant was evaluated by Luminex® Multiplex assay at 7-, 14- and 21-days post-infection. (G) At each time-point IFN-γ/IL-10 ratio was calculated. For each donor, the results obtained were normalized to those of autologous infection. Values for each condition are the mean ± SD of two independent experiments. Means were compared at each time-points using paired t-test.

## DISCUSSION

Infections of patients’ PBMCs with autologous Mtb clinical isolates were associated with lower bacterial loads at early time points and a more robust initial Th1-type cytokine response. As shown in previous studies, Mtb clinical isolates from different lineages exhibited distinct patterns in bacterial load dynamics and dormancy induction (Arbués et al., 2024; Homolka et al., 2010; Mourik et al., 2019; Wang et al., 2010). The restriction of Mtb growth was associated with a stronger production of pro-inflammatory cytokines TNF-α and IFN-γ in the granuloma microenvironment, as Th1 immune response has long been associated with protection against TB infection (Gideon et al., 2022). These findings suggests that an autologous host-Mtb isolate granuloma is of interest as a preclinical tool to further study host regulatory pathways and evaluate therapeutic interventions (Silva-Miranda et al., 2015).

However, this model using patients’ immune cells has limitations. The small blood sample volume restricts cell availability, requiring a miniaturized setup and streamlined analyses. This study highlights the importance of early time points in assessing immune response specificity, particularly in capturing initial T cell responses and broader immune interactions. Refining the model to preserve differences observed at later time points could allow a deeper analysis of autologous host-pathogen pairing. Using cells from TB patients at the start of treatment, rather than the end, might amplify these differences. Besides, future studies incorporating advanced single-cell technologies, such as CITE-seq or high-dimensional cytometry, are needed to explore immune dynamics with greater resolution, offering a deeper understanding of host immune specificity in response to Mtb clinical isolates.

In summary, *in vitro* Mtb granuloma model is useful for investigating host-pathogen specific interactions and lays the foundations for use as a preclinical tool.

## MATERIALS AND METHODS

### Study design

Three patients enrolled in the clinical trial NCT04271397 were included in this study. These patients were all male, aged between 31 and 48 years, originating from Africa, Europe, and Asia respectively. They are referred in this study as P1, P2 and P3, respectively. These patients were selected because they came from varied geographical origins and were infected with clinical Mtb isolates from different lineages—lineages 1, 4, and 3 (L1, L4, and L3), respectively. Additionally, a lineage 2 (L2) Mtb clinical isolate responsible of a large regional cluster (Genestet et al., 2019) and the H37Rv reference strain were included for comparison. Blood samples were collected in heparinized vials at the end of anti-TB treatment for peripheral blood mononuclear cells (PBMCs) isolation by Ficoll-Hypaque (Sigma-Aldrich, Saint-Quentin-Fallavier, France) density-gradient. Aliquots were cryopreserved in fetal bovine serum with 10% DMSO (Sigma-Aldrich) and stored in liquid nitrogen. Between 1.10^7^ et 2.10^7^ PBMC were available per patient.

### Generation of *in vitro* granulomas

PBMCs from TB patients were thawed and washed twice in RPMI with 10% fetal bovine serum. Trypan blue dye exclusion method was used to confirm sample viability greater than 95%. PBMC were adjusted at a density of 10^6^ cells/mL in RPMI supplemented with 10% human AB serum (Sigma-Aldrich) and seeded in 96-well culture plate coated with collagen at 10 µg/cm^2^ (Sigma-Aldrich) and fibronectin at 0.1 µg/cm^2^ (Sigma-Aldrich) for overnight recovery. PBMC were then infected with bacterial suspension in log phase at a multiplicity of infection (MOI) of 1:100 (bacteria to cells) in RPMI supplemented with 10% human AB serum, at 37 °C in 5% CO_2_ atmosphere, to induce spontaneous formation of *in vitro* granulomas. PBMC from each patient were infected with their autologous Mtb clinical isolate as well as with the three other Mtb clinical isolates selected in this study or the H37Rv reference strain. The same bacterial suspension was used to infect PBMCs from all three patients to avoid any bias in the initial bacterial load. Two independent experiments were performed.

### Bacterial growth and dormancy assessment

At 7-, 14- and 21-days post-infection, all well contents were collected, and cells were recovered after coating digestion by a 15 min incubation at 37°C in the presence of collagenase 1 mg/mL (Thermo Fisher Scientific Inc, Rockford, IL, USA). Recovered cells were lysed with distilled water containing 0.1% Triton X100 (Sigma-Aldrich) for 10 min, followed by 3 min of vortex agitation in presence of 1-mm diameter glass beads to disrupt bacterial clumps.

To assess bacterial growth dynamics, mycobacteria were plated in duplicate on 7H10 agar supplemented with 10% OADC (Oleic acid, Albumin, Dextrose, Catalase; Becton Dickinson, Sparks, MD) and incubated for 3-4 weeks at 37°C. To assess bacterial dormancy, mycobacteria were inoculated in MGIT (Mycobacteria Growth Indicator Tube) for which a quarter of the medium volume was replaced or not by autologous Mtb culture supernatant in stationary phase containing resuscitation promoting factors (RPF) (Peters et al., 2023). Analysis of the fluorescence was used to determine if the tube was instrument positive; i.e., the test sample contains viable organisms and results were expressed using MGIT time to positivity (TTP) system, reflecting the bacterial growth (Bowness et al., 2015; Genestet et al., 2021). Quantification of Mtb dormancy was assessed by analyzing the delta between TTP without and with RPF.

### Cytokine release assay

At each time-point, cell supernatant was recovered, and cytokines release was assessed using Bio-Plex Pro Human Cytokine Th1/Th2 Assay (Bio-Rad Laboratory, Hercules, USA) on a FLEXMAP 3D^®^ analyzer (Luminex, Austin, USA). Data were analyzed using Bio-Plex Manager software version 6.1 (Bio-Rad Laboratory). Due to inter-donor variability in cytokine production, each donor’s individual response to a given Mtb isolate was normalized by dividing the response by the average cytokine level across all Mtb isolates for that same donor, enabling a standardized comparison across donors.

### Statistical analysis

Statistical analyses were performed with Graph Pad Prism 5. Values were expressed as the mean ± standard deviation (SD) of two independent experiments. Means were compared using Repeated Measures ANOVA test and Two-Way ANOVA followed by Bonferroni test as specified in Figure legends. \**p*<0.05, ** *p* <0.01, *** *p* <0.001.

## TRANSPARENCY DECLARATIONS

The authors declare that they have no known competing financial interests or personal relationships that could have appeared to influence the work reported in this paper.

## SUPPLEMENTARY FIGURES

**Figure S1:**
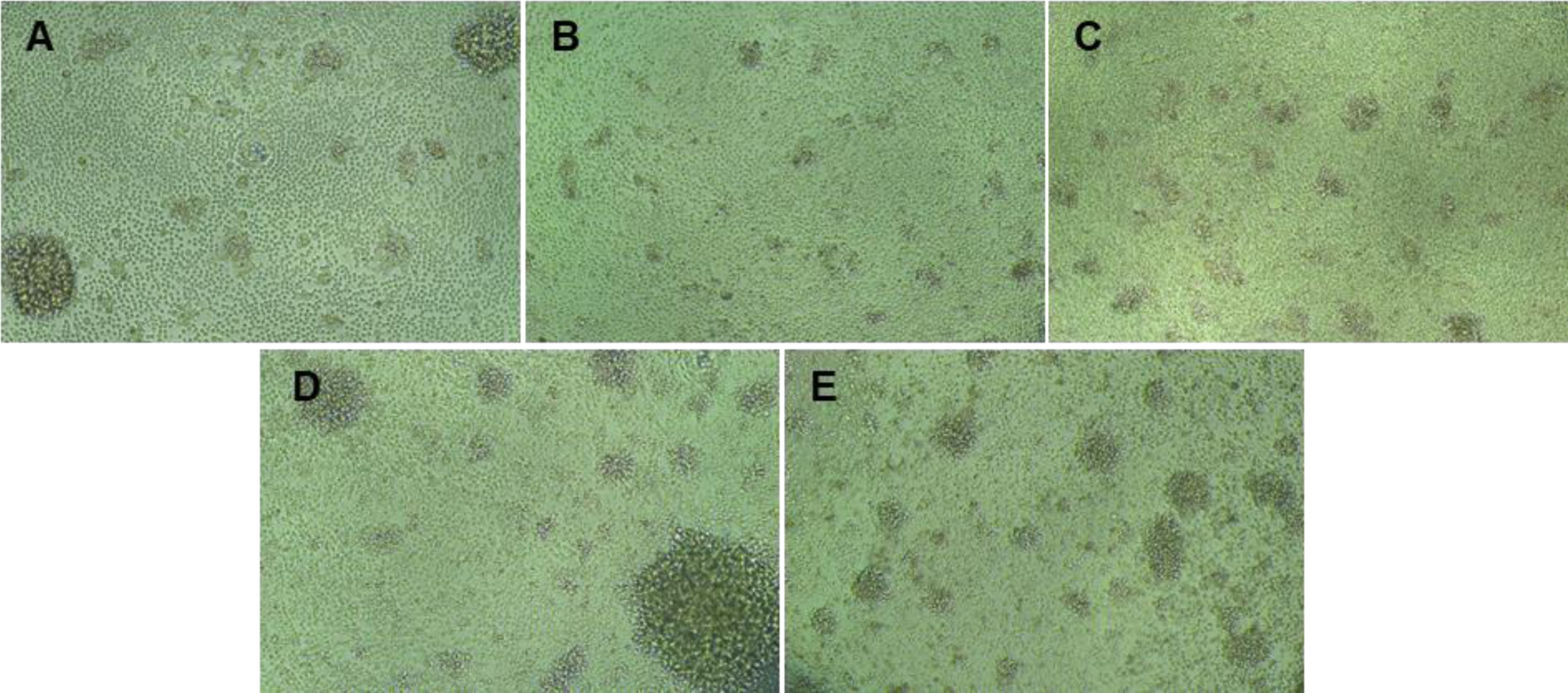
Granuloma formation after PBMC infection by Mtb clinical isolates or H37Rv Mtb reference strain. PBMCs from active TB patient were collected at the end of TB treatment. For *in vitro* granuloma formation, these PBMCs were infected with Mtb clinical isolates belong to the lineage 1 (A), 2 (B), 3 (C), 4 (D) or the H37Rv Mtb reference strain (E). Representative images of granuloma formed upon infection at 14-days post-infection.

**Figure S2:**
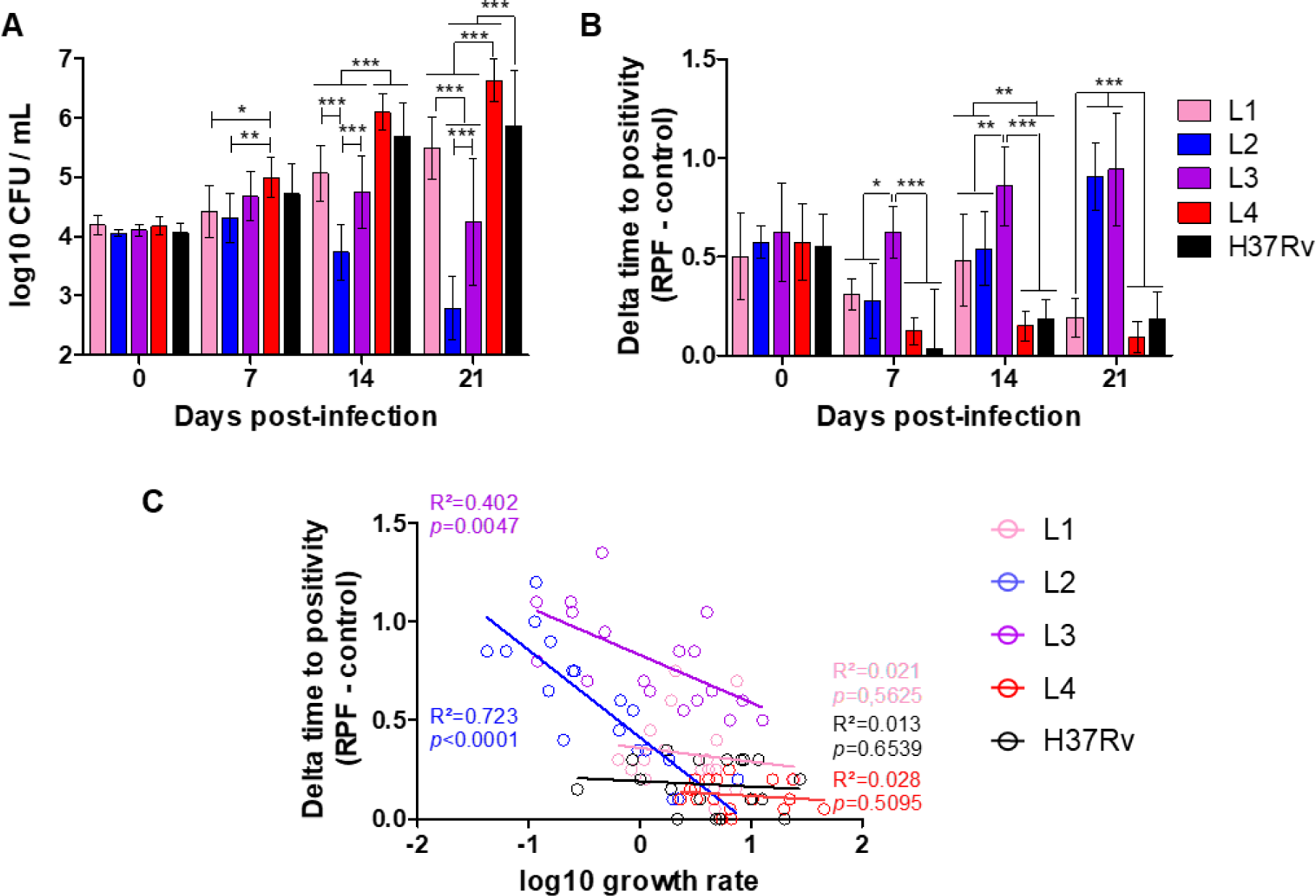
Mtb isolate-dependent bacterial growth and dormancy dynamics. PBMC from 3 active TB patients were collected at the end of anti-TB treatment. For *in vitro* granuloma formation, these PBMCs were infected with their own autologous Mtb clinical isolate as well as with the Mtb clinical isolates from the other two patients, belong to the lineage 1 (pink), lineage 3 (purple) and lineage 4 (red). Additionally, a lineage 2 [L2 (blue)] Mtb clinical isolate from a large regional cluster and the H37Rv reference strain (black) were included for comparison. (A) Bacterial load was assessed by CFU counting after granuloma disruption and cell lysis at 7-, 14-, and 21-days post-infection. (B) At the same time-points, Mtb dormancy was evaluated by quantifying differentially culturable bacteria in the presence or absence of resuscitation-promoting factors (RPF) using MGIT time to positivity (TTP) system, reflecting the bacterial growth. Values for each condition are the mean ± SD of two independent experiments, performed in duplicate. Means were compared using Two-Way ANOVA followed by Bonferroni test. (C) Correlation between Mtb growth rate between time points and Mtb dormancy was evaluated for each Mtb clinical isolate and for the H37Rv reference strain.

**Figure S3:**
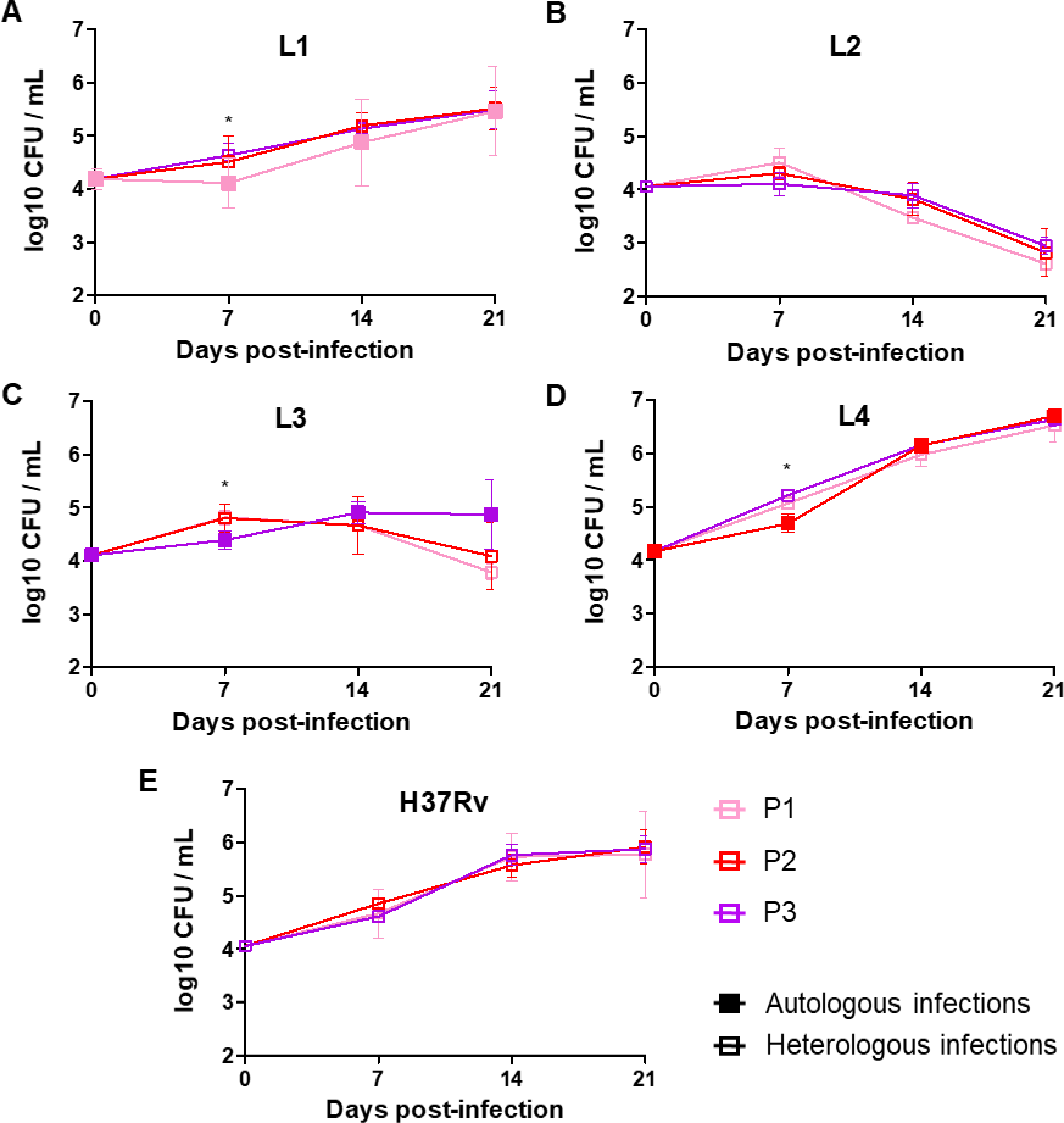
Improved Early Infection Control in Autologous Pairings. PBMC from 3 active TB patients [P1 (pink square), P2 (red square) and P3 (purple square)] were collected at the end of TB treatment. Each patient was infected with a clinical Mtb isolate from a different lineage—lineages 1, 4, and 3 (L1, L4 and L3), respectively. For *in vitro* granuloma formation, these PBMCs were infected with their own autologous Mtb isolate (full square) as well as with the isolates from the other two patients (empty square). Additionally, a lineage 2 (L2) clinical isolate from a large regional cluster and the H37Rv reference strain were included for comparison. Bacterial load was assessed by CFU counting after granuloma disruption and cell lysis at 7-, 14-, and 21-days post-infection. Values for each condition are the mean ± SD of two independent experiments performed in duplicate. Means were compared at each time-points using Repeated Measures ANOVA test.

**Figure S4:**
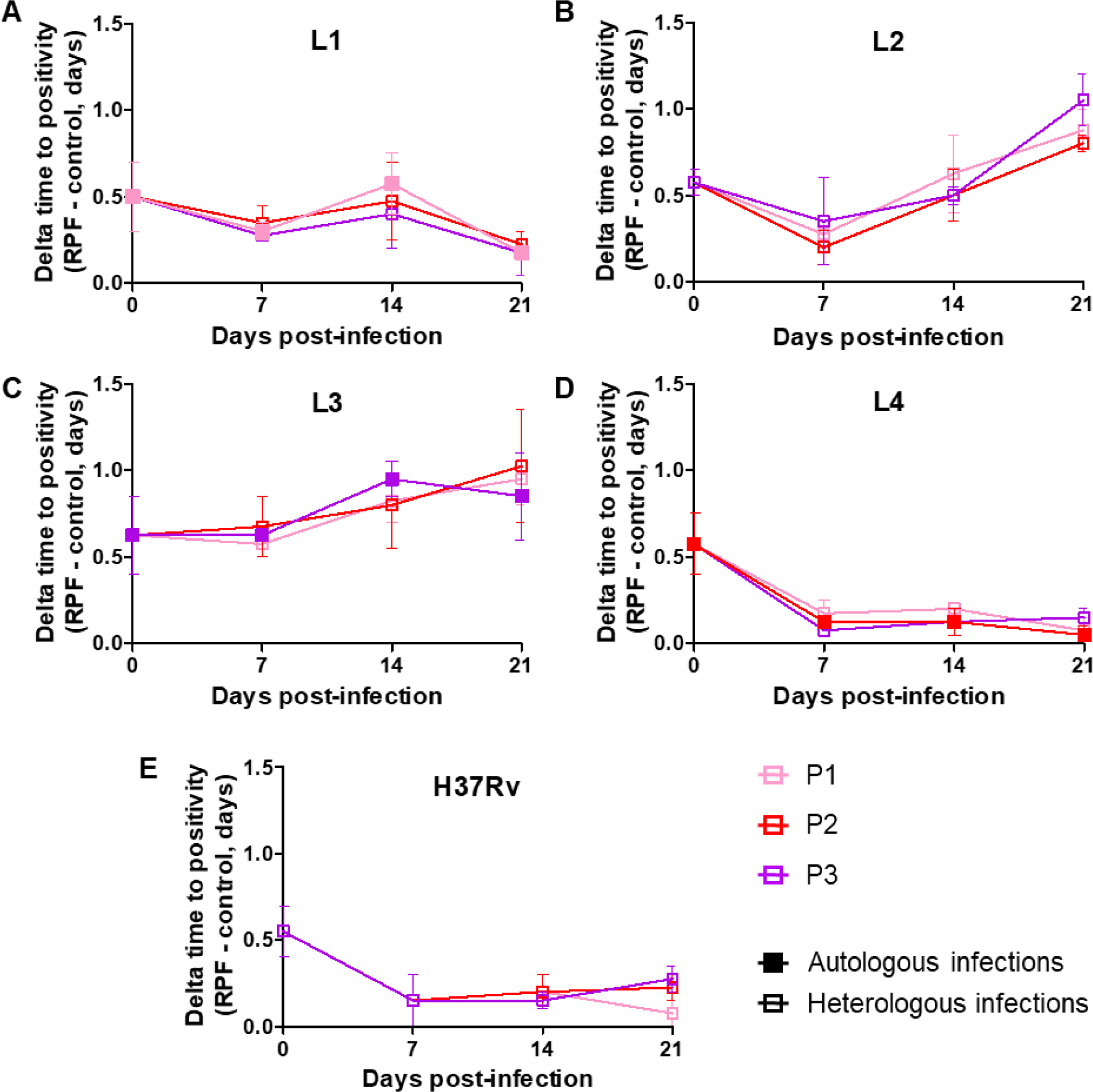
Mtb dormancy was not donor-dependent. PBMC from 3 active TB patients [P1 (pink square), P2 (red square) and P3 (purple square)] were collected at the end of anti-TB treatment. Each patient was infected with a Mtb clinical isolate from a different lineage—lineages 1, 4, and 3 (L1, L4 and L3), respectively. For *in vitro* granuloma formation, these PBMCs were infected with their own autologous Mtb clinical isolate (full square) as well as with the Mtb clinical isolates from the other two patients (empty square). Additionally, a lineage 2 (L2) Mtb clinical isolate from a large regional cluster and the H37Rv reference strain were included for comparison. At 7-, 14-, and 21-days post-infection, Mtb dormancy was evaluated by quantifying differentially culturable bacteria in the presence or absence of resuscitation-promoting factors (RPF) using MGIT time to positivity (TTP) system, reflecting the bacterial growth. Values for each condition are the mean ± SD of two independent experiments. Means were compared at each time-points using Repeated Measures ANOVA test.

**Figure S5:**
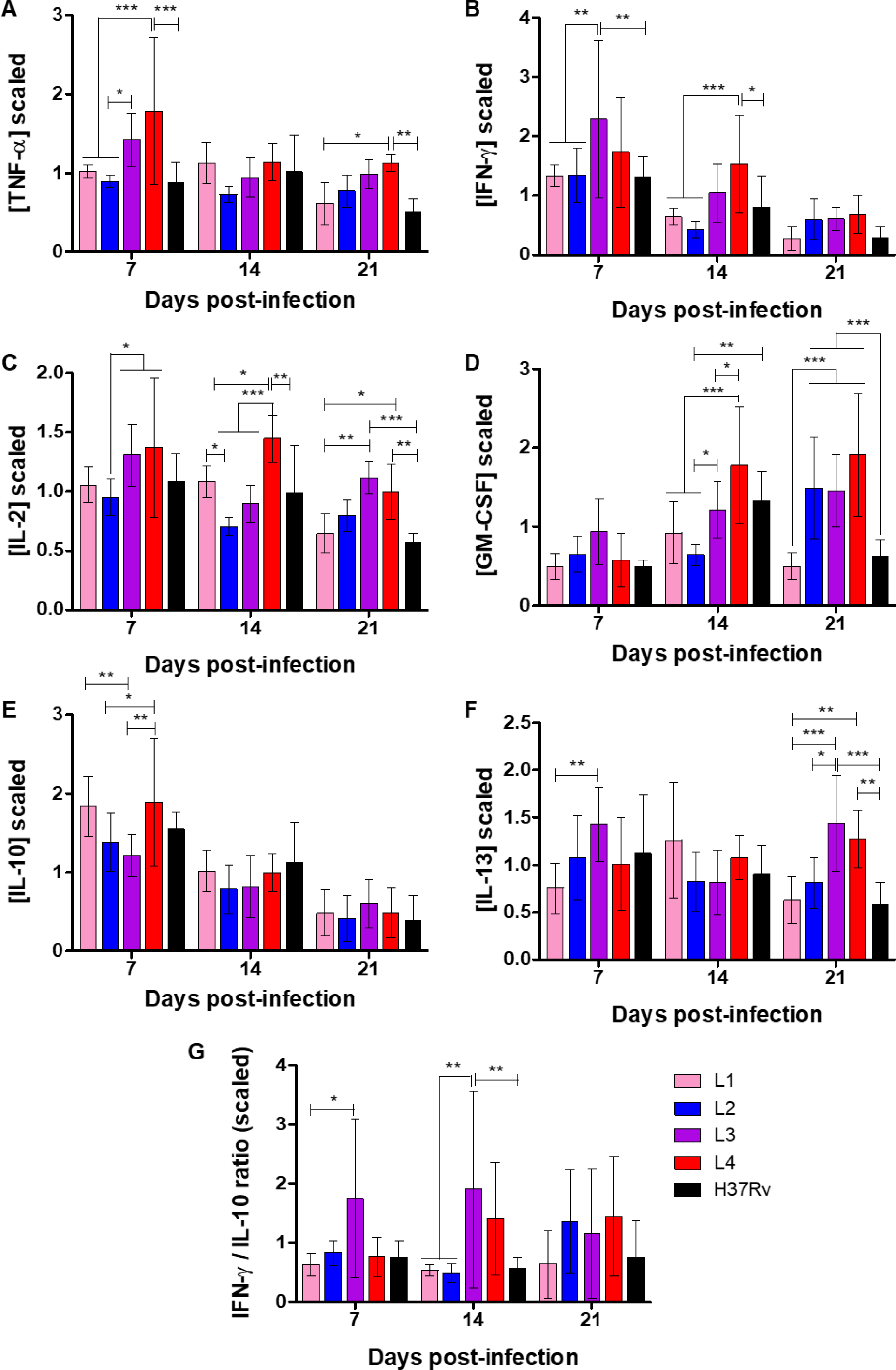
Strain-dependent cytokine production dynamics during Mtb infection. PBMC from 3 active TB patients were collected at the end of anti-TB treatment. For *in vitro* granuloma formation, these PBMCs were infected with their own autologous Mtb clinical isolate as well as with the Mtb clinical isolates from the other two patients, belong to the lineage 1 (pink), lineage 3 (purple) and lineage 4 (red). Additionally, a lineage 2 [L2 (blue)] Mtb clinical isolate from a large regional cluster and the H37Rv reference strain (black) were included for comparison. (A) TNF-α, (B) IFN-γ, (C) IL-2, (D), GM-CSF (E) IL-10, (F) IL-13 release in cell culture supernatant was evaluated by Luminex® Multiplex assay at 7-, 14- and 21-days post-infection. (G) At each time-point IFN-γ/IL-10 ratio was calculated. Values for each condition are the mean ± SD of two independent experiments. Means were compared using Two-Way ANOVA followed by Bonferroni test.

**Figure S6:**
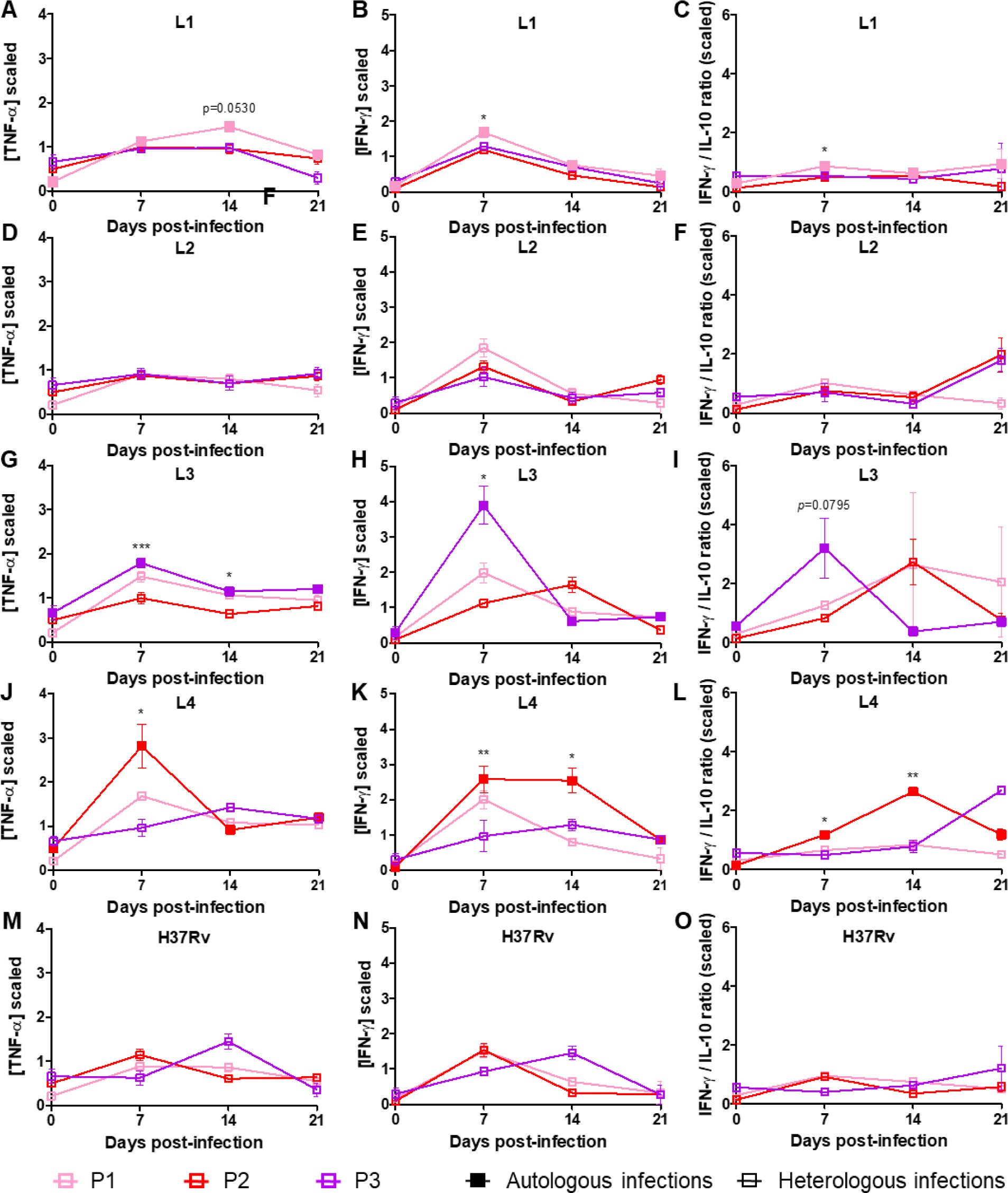
Enhanced early pro-inflammatory response in autologous infections. PBMC from 3 active TB patients [P1 (pink square), P2 (red square) and P3 (purple square)] were collected at the end of TB treatment. Each patient was infected with a clinical Mtb isolate from a different lineage—lineages 1, 4, and 3 [L1 (A-C), L4 (J-L) and L3 (G-I)], respectively. For *in vitro* granuloma formation, these PBMCs were infected with their own autologous Mtb isolate (full square) as well as with the isolates from the other two patients (empty square). Additionally, a lineage 2 [L2 (D-F)] clinical isolate from a large regional cluster and the H37Rv reference strain (M-O) were included for comparison. (A, D, G, J, M) TNF-α, (B, E, H, K, N) IFN-γ, and IL-10 release in cell culture supernatant was evaluated by Luminex® Multiplex assay at 7-, 14- and 21-days post-infection. (C, F, I, L, O) At each time-point IFN-γ/IL-10 ratio was calculated. Values for each condition are the mean ± SD of two independent experiments. Means were compared at each time-points using Repeated Measures ANOVA test.

**Figure S7:**
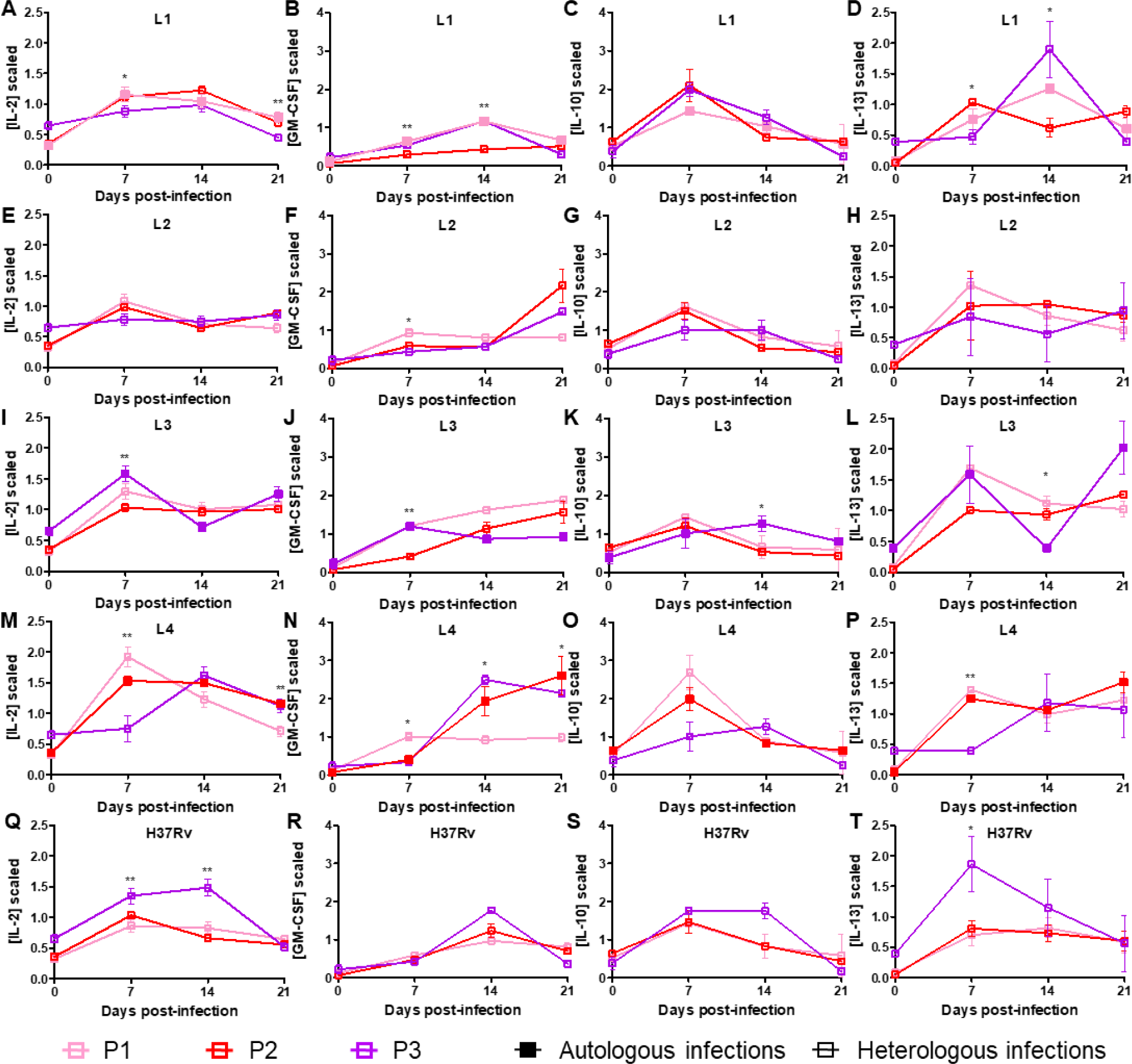
Cytokine production dynamics in autologous and heterologous infections. PBMC from 3 active TB patients [P1 (pink square), P2 (red square) and P3 (purple square)] were collected at the end of anti-TB treatment. Each patient was infected with a Mtb clinical isolate from a different lineage—lineages 1, 4, and 3 [L1 (A-D), L4 (M-P) and L3 (I-L)], respectively. For *in vitro* granuloma formation, these PBMCs were infected with their own autologous Mtb clinical isolate (full square) as well as with the Mtb clinical isolates from the other two patients (empty square). Additionally, a lineage 2 [L2 (E-H)] Mtb clinical isolate from a large regional cluster and the H37Rv reference strain (Q-T) were included for comparison. (A, E, I, M, Q) IL-2, (B, F, J, N, R) GM-CSF, (C, G, K, O, S) IL-10 and (D, H, L, P, T) IL-13 release in cell culture supernatant was evaluated by Luminex® Multiplex assay at 7-, 14- and 21-days post-infection. Values for each condition are the mean ± SD of two independent experiments. Means were compared at each time-points using Repeated Measures ANOVA test.

**Figure S8:**
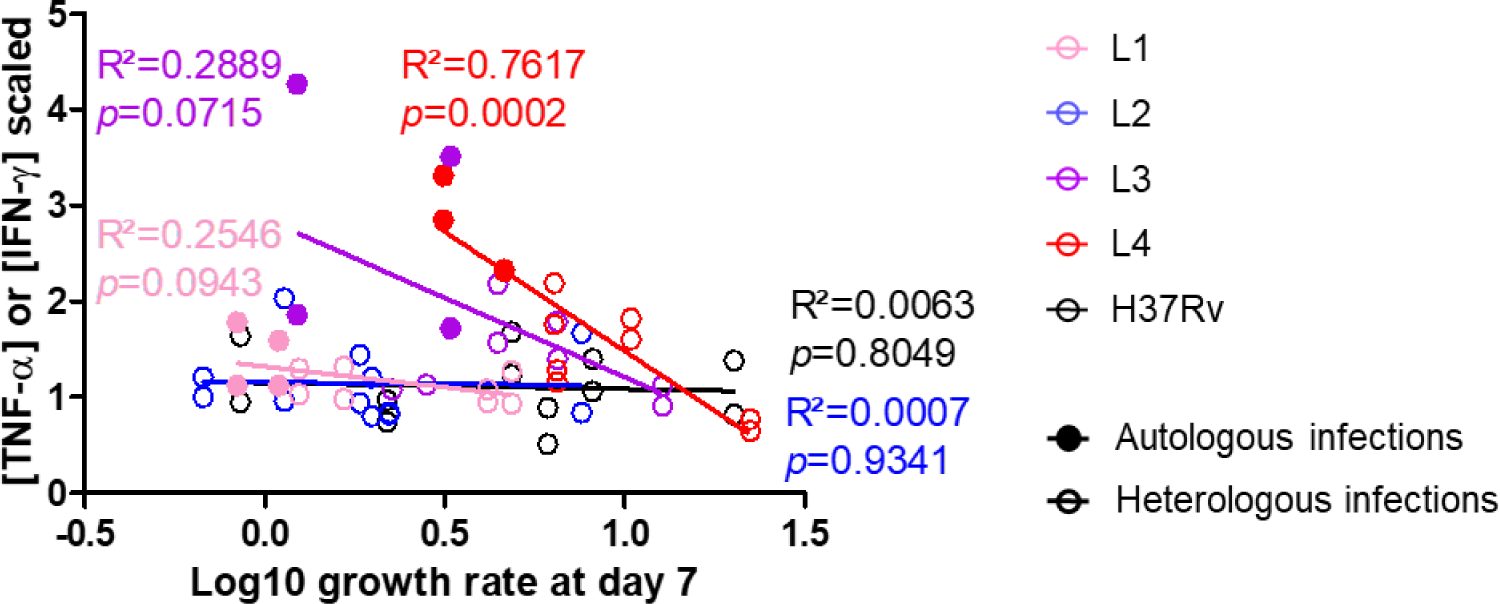
Early bacterial growth rate was inversely correlated with the pro-inflammatory response only in comparisons between autologous and heterologous infections. PBMC from 3 active TB patients were collected at the end of anti-TB treatment. For *in vitro* granuloma formation, these PBMCs were infected with their own autologous Mtb clinical isolate (full circle) as well as with the Mtb clinical isolates from the other two patients (empty circle), belong to the lineage 1 (pink), lineage 3 (purple) and lineage 4 (red). Additionally, a lineage 2 [L2 (blue)] Mtb clinical isolate from a large regional cluster and the H37Rv reference strain (black) were included for comparison. Correlation between Mtb growth rate and TNF-α or IFN-α production at 7-days post-infection was evaluated.

